# Reversion of a RND transporter pseudogene uncovers latent stress resistance in *Brucella ovis*

**DOI:** 10.1101/2025.05.10.653276

**Authors:** Thomas Kim, Bongjin Hong, Thomas V. O’Halloran, Erika Lisabeth, Richard R. Neubig, Aretha Fiebig, Sean Crosson

**Author notes:** Correspondence should be addressed to A.F. or S.C.

## Abstract

Small-molecule screens can advance therapeutic discovery while yielding new insights into pathogen biology. Through a luminescence-based screen, we identified clinically approved dihydropyridines that reduced the fitness of the intracellular pathogen *Brucella ovis* within mammalian phagocytes. Given the established role of dihydropyridines as inhibitors of mammalian L-type calcium channels and our observation that drug treatment perturbed calcium and manganese levels in host phagocytes, we initially hypothesized a host-directed mechanism of action. However, dose-response assays in axenic medium revealed that these drugs can directly inhibit *B. ovis* growth. To investigate the genetic basis of *B. ovis* susceptibility to dihydropyridine treatment, we selected for mutants capable of growing in the presence of cilnidipine. Cilnidipine-resistant isolates carried single-base deletions in the *bepE* pseudogene that restored an open reading frame encoding an RND-family transporter subunit. *B. ovis* is an ovine venereal pathogen that has experienced significant pseudogenization in its recent evolutionary history. Frameshift mutations that restored *bepE* function in *B. ovis* increased its resistance not only to dihydropyridines but also to a broad range of membrane-disrupting agents, including bile acid. Conversely, deleting *bepE* in *Brucella abortus*, a related zoonotic species that retains an intact version of the gene, increased its sensitivity to bile acid in vitro and to cilnidipine in the intracellular niche. We conclude that *bepE* is a key determinant of chemical stress resistance in *Brucella* spp., and that its pseudogenization in *B. ovis* contributes to the documented hypersensitivity of this host-restricted lineage to chemical stressors.

**Author Summary:** *Brucella* species are intracellular bacterial pathogens that infect diverse wildlife and livestock and can also cause disease in humans. We developed a high-throughput assay to identify compounds that reduce the intracellular fitness of *Brucella ovis*, a globally distributed pathogen of sheep. This screen identified two dihydropyridine-class hypertension drugs as inhibitors of *B. ovis* fitness within mammalian cells. While dihydropyridines are established inhibitors of calcium channels in mammals, our results revealed that they also directly inhibited *B. ovis* growth. The *B. ovis* genome contains over 300 pseudogenes, and we discovered that resistance to dihydropyridines and other chemical stressors can emerge through frameshift mutations that resurrect the function of *bepE*, a conserved pseudogene encoding a membrane transporter subunit. In other *Brucella* species, *bepE* remains intact, and we show that deleting this gene in the zoonotic pathogen *Brucella abortus* increases susceptibility to both dihydropyridine treatment and membrane-disrupting stress. Our study provides evidence that pseudogenes like *bepE* can serve as a latent reservoir of adaptive functions in *B. ovis*, including drug and stress resistance.

## Introduction

Brucellosis, a zoonotic disease caused by *Brucella* species, poses a significant threat to human and animal health while also imposing a substantial economic burden on affected regions [1]. Despite vaccination efforts, livestock in many parts of the world remain vulnerable to Brucella infection, leading to reproductive losses, reduced productivity, and trade restrictions [2, 3]. A recent analysis of global public health data conservatively estimated an annual incidence of 2.1 million cases worldwide in humans [4], with infection capable of causing severe complications such as endocarditis, hepatic failure, osteomyelitis, and neurobrucellosis [5]. While antibiotic treatment is available, it requires prolonged combination therapy and is associated with high relapse rates [6, 7] and treatment failure in some cases [8]. The widespread use of antimicrobials in agriculture and human medicine may be driving the emergence of *Brucella* strains with antimicrobial resistance, posing a future threat to effective treatment and disease control [9, 10].

As intracellular pathogens, *Brucella* species subvert host cellular processes to enable their replication, survival, and egress, primarily through the activity of type IV secreted effector proteins [11]. While several host pathways [11, 12] and cellular factors [13] have been identified as contributors to *Brucella* infection and dissemination, much remains to be learned about *Brucella* infection biology. Unbiased small-molecule screens that disrupt host cell functions provide a powerful approach to identify host pathways and proteins that support *Brucella* survival in mammalian cells. These screens may also yield potential therapeutic candidates, including both host-directed and pathogen-targeting molecules for treating brucellosis and other intracellular infections [14].

A previous screen from our group identified small molecules that inhibit *Brucella abortus* growth in axenic cultures, as well as compounds that selectively impair *B. abortus* replication within macrophages while having minimal effects on bacterial growth in axenic conditions [15, 16]. In this study, we developed a luminescence-based screen to extend this work and identify small molecules that inhibit the replication of *Brucella ovis* in THP-1 macrophage-like cells. *B. ovis* is an ovine venereal pathogen with distinct cellular and physiological characteristics that set it apart from *other Brucella* species [17-21], including the absence of O-polysaccharides on its outer membrane [22, 23]. The goal of this work was to identify compounds that inhibit *Brucella* replication in its intracellular niche, and to expand our knowledge of genes that are important for intracellular fitness across *Brucella* species.

Our small molecule screen identified multiple classes of molecules that impair *B. ovis* growth in mammalian cells, including the dihydropyridine-class Ca^2+^ channel blockers nicardipine and cilnidipine, which significantly inhibit *B. ovis* fitness within a host. Here we focus on the genetic determinants of *Brucella* susceptibility to dihydropyridines and explore how pseudogenization in *B. ovis* alters its resistance to diverse chemical stressors. We identify *bepE*, which encodes an inner membrane permease subunit of a tripartite RND-family transporter, as a key factor in dihydropyridine tolerance and membrane stress resistance. While *bepE* is a frameshifted pseudogene in *B. ovis*, mutations that restore its reading frame are easily selected and enhance resistance to multiple chemical stressors, including dihydropyridines. In *B. abortus*, where the *bepE* gene remains intact, its deletion increases sensitivity to bile acids and its susceptibility to cilnidipine treatment in host cells, reinforcing its role as a broad-spectrum chemical stress resistance factor. This study demonstrates how *bepE* contributes to chemical resistance in the *Brucella* species and provides new insight into genes that impact *Brucella* resistance to drug treatment in an intracellular context.

## Results

### Identification of small molecules that inhibit B. ovis replication in THP-1 cells

We screened 1,280 small molecules from the Prestwick Chemical Library to identify compounds that inhibit *B. ovis* replication in mammalian cells. Our initial aim was to identify molecules that impair the fitness of *B. ovis* within the host intracellular niche without exerting direct antibacterial effects in axenic culture. We performed this screen using a luciferase-expressing strain of *B. ovis* ATCC 25840, in which the lux operon was integrated into a neutral chromosomal site. Luminescence served as a proxy for intracellular replication within host cells. This effort yielded 49 compound hits that met our criteria for selectively inhibiting *B. ovis* in the intracellular niche (Table S1). To identify potential host cellular pathways that influence *B. ovis* fitness in THP-1 macrophages, we prioritized compounds with well-characterized mechanisms of action, including several FDA-approved drugs. Notably, multiple hits affected calcium (Ca^2+^) homeostasis or dopamine signaling (Table S1). We chose 10 compounds for validation with chemical stocks used for the initial screening, assessing their cytotoxicity and their potency in the context of intracellular and axenic growth conditions. Dose-response assays confirmed four of these—namely nicardipine, cilnidipine, lomerizine, and bifonazole—as inhibitors of intracellular replication that were not toxic to host cells at the assessed concentration (Table 1; Fig. S1).

**Table 1.** Validation of select hits from the Prestwick Chemical Library. The axenic IC_50_, intracellular IC_50_ and the CC_50_ values were calculated from dose-response studies using stocks from the chemical library. Hits considered *bona fide* host targeted molecules from these validation assays are designated with an asterisk.

| Compound name | Therapeutic class | Major pathways | Intracellular $\text{IC}_{50}$ ($\mu\text{M}$) | Axenic $\text{IC}_{50}$ ($\mu\text{M}$) | Cytotoxic $\text{CC}_{50}$ ($\mu\text{M}$) |
| --- | --- | --- | --- | --- | --- |
| Nicardipine* | Antihypertension | Ca | 2 | >50 | 49 |
| Cilnidipine* | Antihypertension | Ca | 2 | 5 | >50 |
| Lomerizine* | Antimigraine | Ca | 9 | >50 | 50 |
| Aripiprazole | Antipsychotic | GPCR/Dopamine | 4 | >50 | 12 |
| Bifonazole* | Antifungal | Cytochrome P450 | 14 | 4 | >50 |
| Dipyridamole | Antithrombotic | Phosphodiesterase | >50 | >50 | 41 |
| Racecadotril | Antidiarrheal | Opioid | >50 | >50 | >50 |
| Phenacetin | Analgesic | Prostaglandin | >50 | >50 | >50 |
| Tranylcypromine | Antidepressant | Monoamine oxidase | >50 | >50 | >50 |
| Halofantrine | Antimalarial | Hematin binding | >50 | 11 | >50 |

Previous screens identified calcium channel blockers as inhibitors of *B. abortus* replication within the intracellular niche [15, 16], and a recent case–control study concluded that the use of dihydropyridine-type Ca^2+^ channel blockers lowered the risk of active tuberculosis [24]. For tuberculosis, the effect of these compounds on disease development may relate to the role of L-type Ca^2+^ channels in supporting iron import [25], which is crucial for *Mycobacterium tuberculosis* fitness in host cells. Given the clinical potential of Ca^2+^ channel blockers for the treatment of intracellular infections and their emergence in our drug screens against *Brucella* growth, we prioritized two FDA-approved dihydropyridine-class Ca^2+^ channel blockers—nicardipine and cilnidipine— for further investigation. Both are widely used to treat hypertension and angina pectoris [26].

### Differential activity of dihydropyridines against Brucella in host cells and axenic culture

We next re-evaluated the effects of nicardipine and cilnidipine in our assays using freshly prepared stock solutions to more quantitatively establish potency. Both drugs showed a similar potency for inhibiting intracellular growth of *B. ovis*, with comparable IC50 values of 2.3 and 2.5 µM, respectively, when growth was assessed after 48 h of treatment (Fig. 1A). To assess whether this inhibition was due to an effect on the host cell or *Brucella* cells, we also conducted dose-response assays in static axenic *B. ovis* cultures, i.e. in the absence of THP-1 cells (Fig. 1B). Although the two drugs are structurally similar (Fig. 1C), cilnidipine inhibited *B. ovis* growth in a complex liquid broth at concentrations above 5 µM, while nicardipine did not affect the terminal density of *B. ovis* at any concentration tested (Fig. 1B).

**Figure 1.**
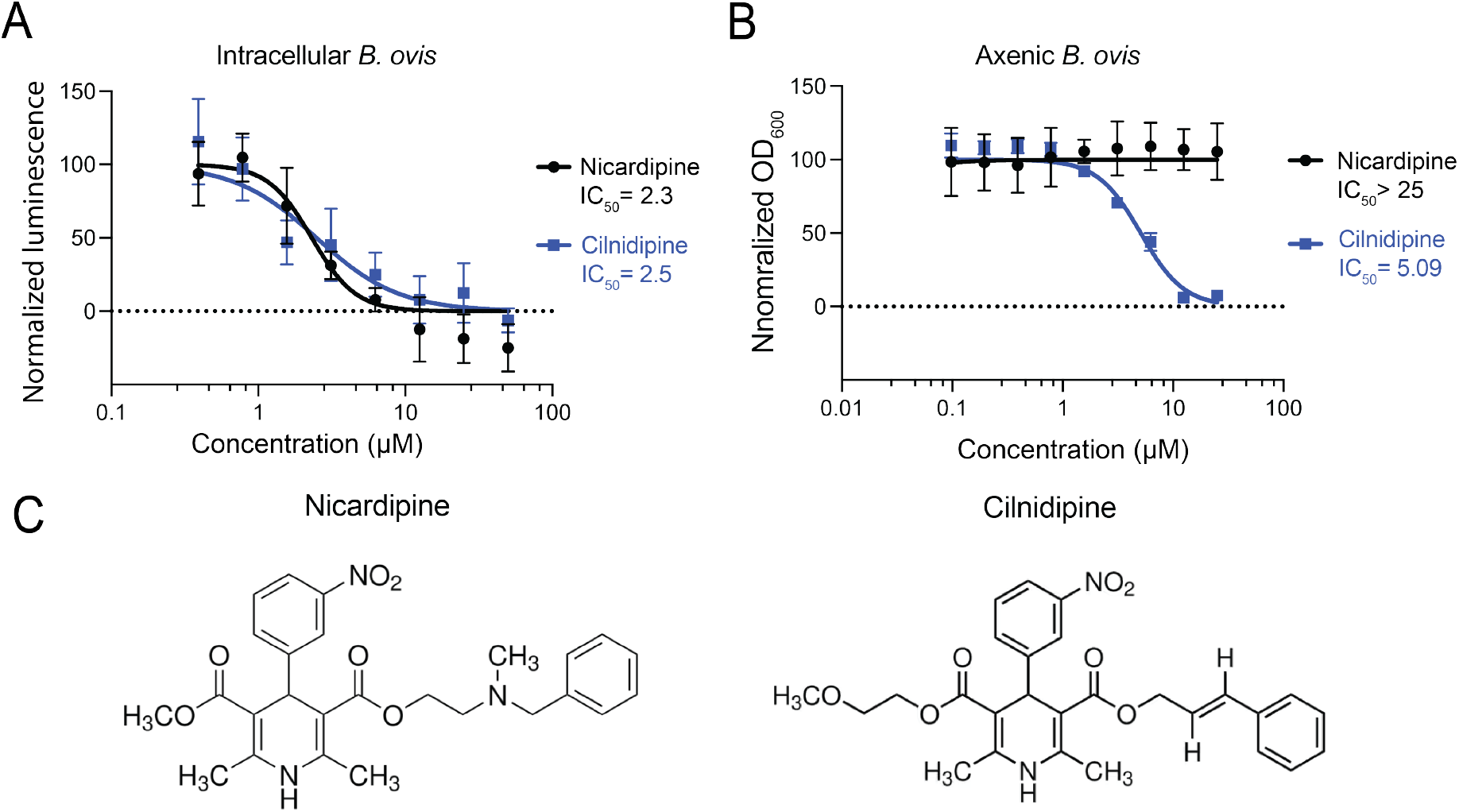
The dihydropyridine Ca^2+^ blockers nicardipine and cilnidipine inhibit intracellular *B. ovis* growth and cilnidipine inhibits *B. ovis* axenic growth. **(A)** Intracellular inhibitory activities of nicardipine and cilnidipine during *B. ovis* THP-1 macrophage infection. *B. ovis* luminescence was measured 48 h post-treatment and normalized to signal from infected untreated cells. IC50 values were calculated using GraphPad Prism software. **(B)** Axenic inhibitory activity of nicardipine and cilnidipine during *B. ovis* growth in liquid medium. Optical densities at 600 nm were measured after 48 h of growth and normalized to untreated cultures. **(C)** Chemical structures of nicardipine and cilnidipine. In A and B, data represent the mean and standard deviation of 3 biological replicates.

### Nicardipine treatment alters the THP-1 metallome

Nicardipine and cilnidipine are well-known inhibitors of Ca^2+^ import into vascular smooth muscle and cardiac cells [26] but, to our knowledge, their effect on levels of Ca^2+^ or other metals in monocytic cell lines such as THP-1 has not been examined. To evaluate intracellular metal levels following dihydropyridine treatment, we used triple quadrupole inductively coupled plasma mass spectrometry (ICP-QQQ) to analyze a panel of elements in THP-1 macrophage-like cells (Fig. 2, Fig. S2). The cells were treated with 25 µM nicardipine, a concentration near the reported IC50 for L-type Ca^2+^ channels at membrane potentials of −15 to −30 mV [27], a range typical of human macrophages and monocytic cells [28, 29]. Metal concentrations were normalized to phosphorus, of which advantages for intracellular metal analysis have been shown in multiple studies [30-33]. Most metal levels remained unchanged following nicardipine treatment (Fig. S2). However, we observed a ∼20–40% increase in intracellular calcium and manganese levels in treated THP-1 cells (Fig. 2A and 2B).

**Figure 2.**
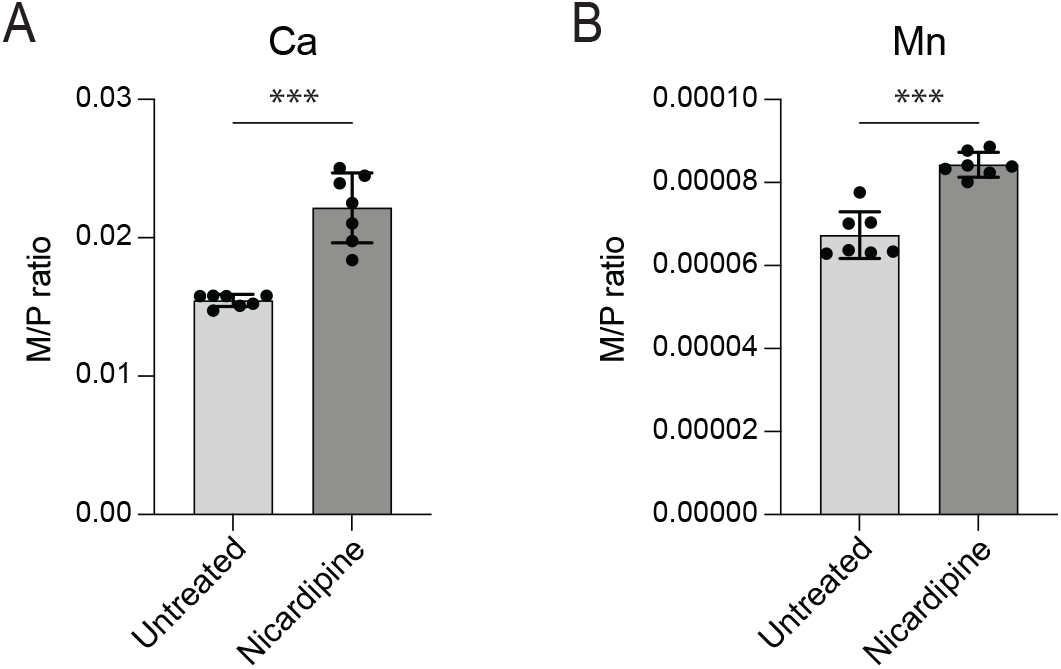
Nicardipine treatment alters the THP-1 macrophage metallome. Elemental content of untreated THP-1 cells or cells treated with 25 µM nicardipine for 48 hours was determined using triple quadrupole inductively coupled plasma mass spectrometry (ICP-QQQ). Each element was normalized by the total phosphorus in each sample (M/P). **(A)** Calcium and **(B)** Manganese levels were significantly elevated upon nicardipine treatment. Other elements are presented in Fig. S2. Bars represent the mean ± standard deviation of 7 biological replicates measured over 2 independent experiments. The full metallome data set was analyzed using multiple unpaired t-tests and Bonferroni-Dunn method to adjust for multiple comparisons (***, adjusted P < 0.001).

The mechanism underlying elevated calcium remains unclear but may reflect compensatory calcium influx triggered by depletion of intracellular Ca^2+^ stores or altered activity of alternate Ca^2+^ transport pathways [34, 35]. Notably, inhibition of voltage-gated Ca^2+^ channels has been shown to promote calcium influx in dendritic cells and reduce *Mycobacterium tuberculosis* burden in mice [36], supporting a model in which nicardipine impacts calcium signaling in immune cells. Increased manganese levels following nicardipine treatment is notable, as elevated intracellular Mn has been linked to activation of host-defense pathways against microbial infection [37, 38]. Together, our results indicate that nicardipine selectively alters intracellular calcium and manganese levels in a human monocytic cell line. Although the downstream effects of these metal shifts on host cell physiology are not yet defined, they may help explain the enhanced efficacy of nicardipine against *B. ovis* in the intracellular niche compared to axenic conditions (Figure 1).

### Mutations in transporter genes confer resistance to dihydropyridines in axenic culture

Nicardipine disrupted calcium and manganese levels in THP-1 host cells (Fig. 2), which may influence *B. ovis* fitness within the intracellular niche. However, cilnidipine exhibited direct toxicity to *B. ovis* in axenic culture, as measured by a terminal density assay (Fig. 1B), suggesting that the intracellular inhibition observed upon dihydropyridine treatment may be at least partly independent of host-mediated effects. We hypothesized that *B. ovis* susceptibility to drug is influenced by specific bacterial genes. To identify such genes, we selected for spontaneous mutants capable of growing on solid medium supplemented with 25 µM cilnidipine and recovered multiple resistant isolates. Whole-genome sequencing revealed mutations in one of two genes: *bepE* (locus BOV_RS01560/BOV_0306) or emrA (BOV_RS10885/BOV_A0108) (Table 2). *bepE* encodes a predicted inner membrane permease subunit of a resistance-nodulation-division (RND) family transporter system, while *emrA* encodes a component of a major facilitator superfamily (MFS) multidrug efflux pump. Mutations in *emrA* resulted in single amino acid substitutions (L271P or A102T), which likely conferred resistance by enhancing efflux activity. *bepE* is one of many genes that have undergone pseudogenization in the *B. ovis* lineage through indel mutations that cause frameshifts [18, 19, 21]. Cilnidipine-resistant isolates carried single-base deletions near the original pseudogene frameshift site that extended the open reading frame (ORF) from 1,689 to 3,156 nucleotides (Fig. 3A–B). The resulting “mutant” *bepE* alleles encode proteins aligning with full-length *bepE* homologs in *B. melitensis, B. abortus, B. suis, B. canis*, and other *Brucella* species (Fig. 3C-D), effectively reconstituting a functional gene. To assess the functional impact of bepE ORF restoration, we compared growth of wild-type (WT) *B. ovis* and a *bepE*-reconstituted mutant (ΔG at position 1,500) in liquid culture with 25 µM nicardipine or cilnidipine. In untreated media, both strains grew similarly (Fig. S3). While a terminal density assay showed no effect of 25 µM nicardipine on *B. ovis* (Fig. 1), a kinetic growth assay revealed a modest lag in the growth of WT cultures treated with nicardipine; the restored *bepE* mutant showed no lag in the presence of nicardipine (Fig. S3). WT cultures failed to grow over 72 h in the presence of 25 µM cilnidipine, whereas the *bepE* mutant resumed growth after a 24 h lag and reached densities comparable to untreated WT by 72 h (Fig. S3). These results demonstrate that nicardipine and cilnidipine can directly inhibit *B. ovis* growth, though to different degrees, and that restoration of the *bepE* pseudogene to encode an open reading frame that matches other *Brucella* species confers drug resistance.

**Table 2.**
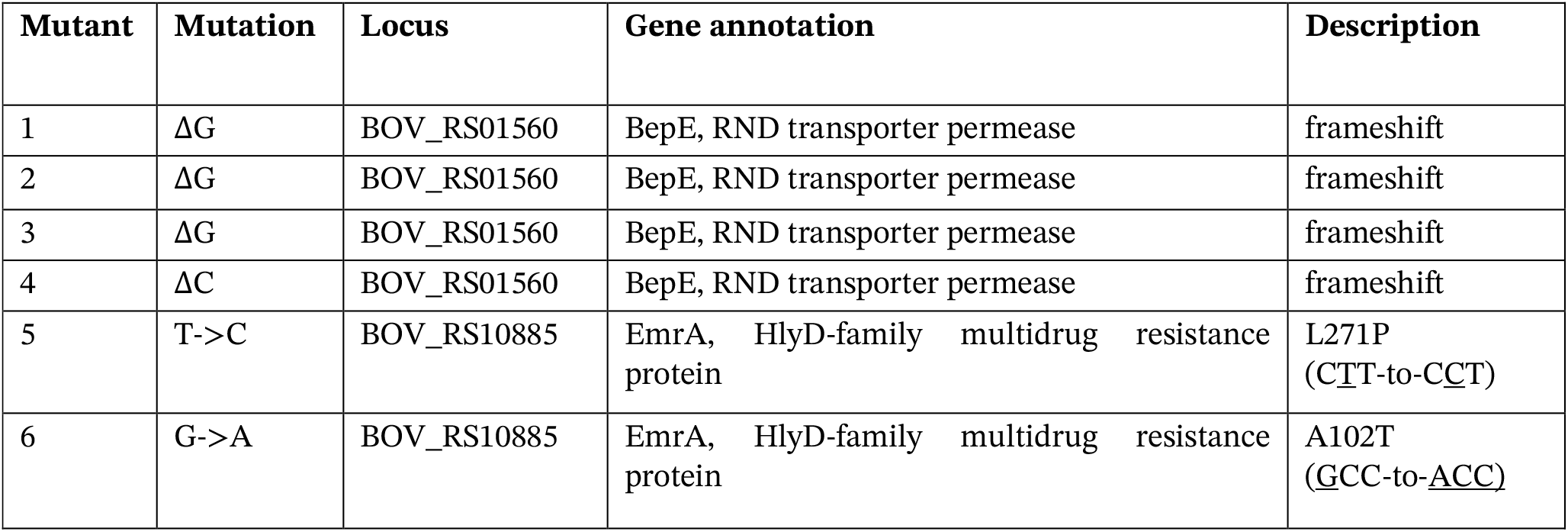
Mutations identified in six independent cilnidipine-resistant spontaneous mutants following growth on solid medium containing 25 µM cilnidipine.

**Figure 3.**
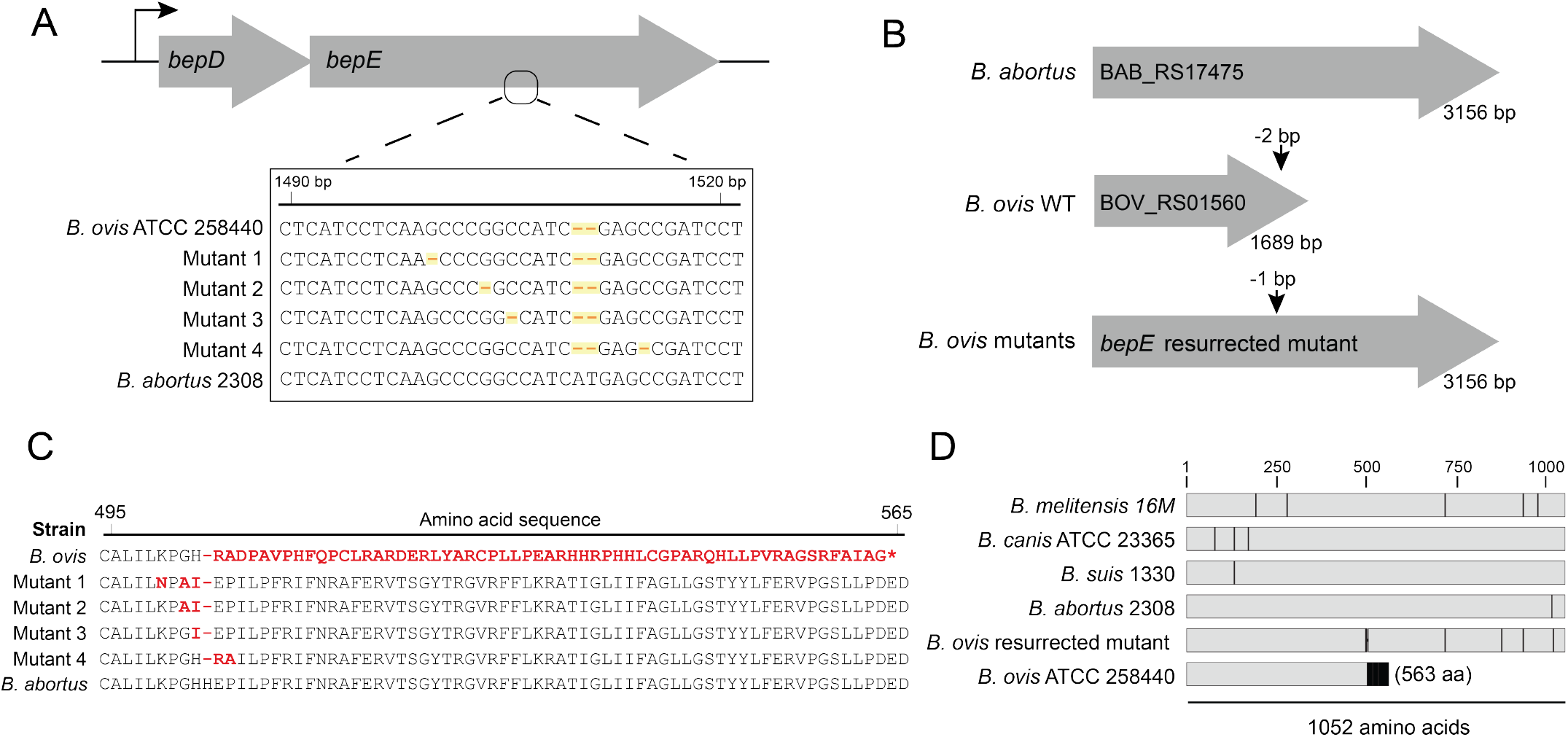
*B. ovis* cilnidipine-resistant mutants harbor single-nucleotide deletions that resurrect the open reading frame of the pseudogene *bepE*. **(A)** Diagram of the *bepDE* locus (top) and alignment of *bepE* nucleotide sequences from wild-type *B. ovis*, four cilnidipine-resistant mutants, and *B. abortus* (bottom). The 2 bp deletion that led to the pseudogenization of *B. ovis bepE*, and the single base deletions in the cilnidipine-resistant mutants are highlighted. **(B)** Schematic of the *bepE* open reading frame *B. abortus* 2308, *B. ovis* ATCC 25840, and a *B. ovis* cilnidipine-resistant mutant. The site of the 2 bp frameshift in wild-type *B. ovis* compared to the other *Brucella* species, and site of the restoring frameshift mutations are indicated. **(C)** Alignment of the partial BepE amino acid from wild-type *B. ovis* ATCC 25840, four cilnidipine-resistant mutants and *B. abortus*. Differences from the consensus of the genus, exemplified by *B. abortus*, are highlighted in red. **(D)** Alignment of the entire BepE protein sequence from representative *Brucella* species lineages and one *B. ovis* cilnidipine-resistant mutant. Differences from the consensus are indicated in black.

### Restoration of the *B. ovis bepE* open reading frame confers broad-spectrum detergent resistance in vitro

*B. ovis* is more sensitive to compounds that disrupt the cell envelope than other *Brucella* species [39, 40], but the genetic basis for this increased sensitivity has not, to our knowledge, been previously described. We hypothesized that the mutations in *bepE* and *emrA* that confer resistance to dihydropyridines might also enhance resistance to other cell envelope stressors. To test this, we evaluated the sensitivity of these mutants to the detergents sodium dodecyl sulfate (SDS) and CHAPS, as well as to the secondary bile acid deoxycholate, which has detergent-like properties. Wild-type (WT) *B. ovis* was highly sensitive to 0.005% (w/v) SDS, 0.009% (w/v) deoxycholate, and 0.005% (w/v) CHAPS. In contrast, *bepE*-restored mutants exhibited resistance to all three compounds at these concentrations (Fig. 4A). Mutations in *emrA* had more limited effects: both *emrA* mutants (L127P and A102T) were as resistant to SDS as the *bepE* mutants and showed improved CHAPS tolerance compared to WT, though to a lesser degree than the *bepE*-restored strains. Only *emrA*(A102T), but not *emrA*(L127P), conferred partial resistance to deoxycholate (Fig. 4A). These results indicate that pseudogenization of *bepE* contributes to the cell envelope stress sensitivity of *B. ovis*.

**Figure 4.**
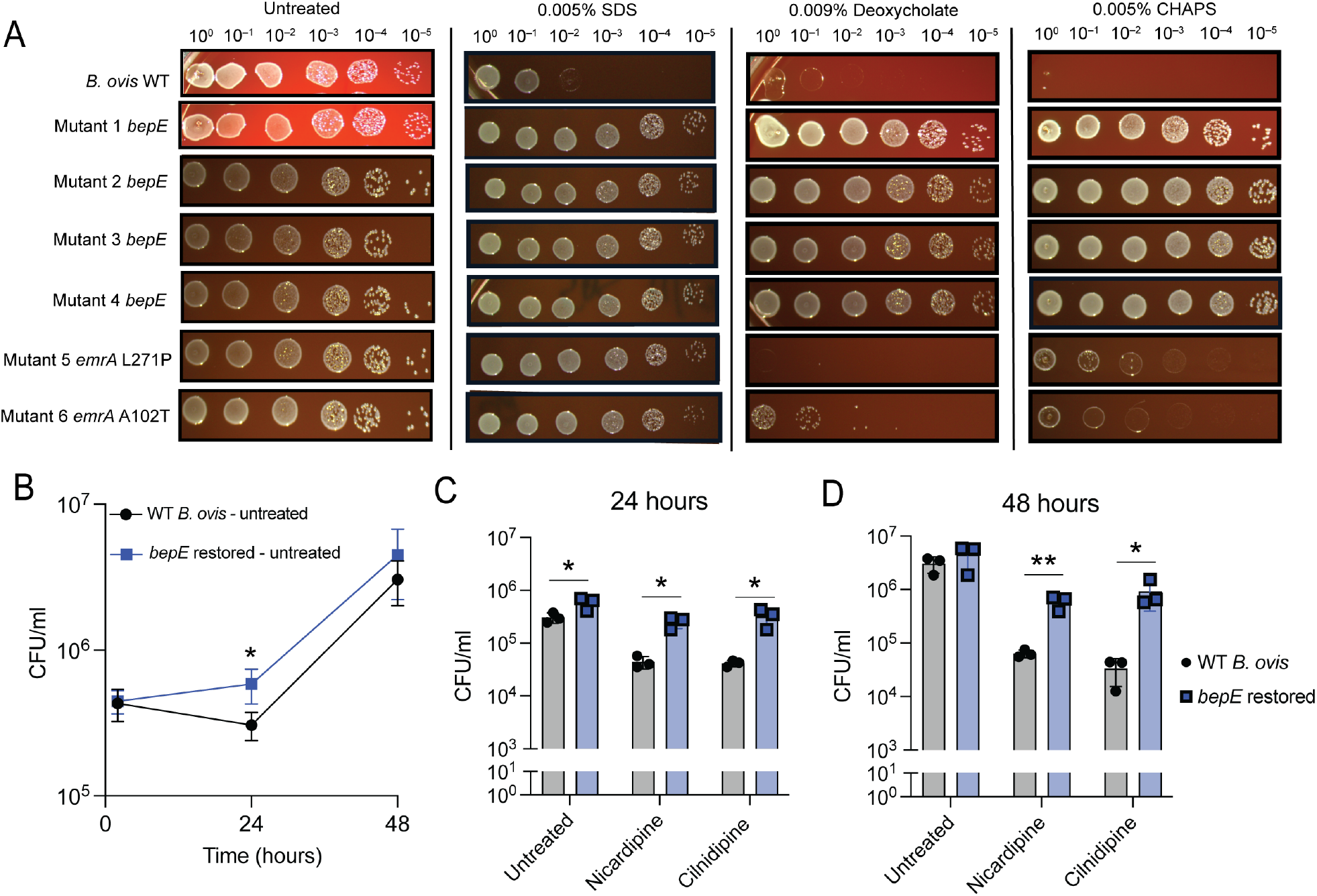
Restoration of *B. ovis* BepE function confers detergent resistance and enhanced tolerance of dihydropyridines during macrophage infection. **(A)** Growth of the wild-type (WT) *B. ovis* and cilnidipine-resistant mutants on TSA blood plates with or without detergents. Cultures of each genotype were normalized by optical densitiy, 10-fold serially diluted, and spotted onto TSA blood plates that were untreated or contained 0.005% SDS, 0.009% deoxycholate, or 0.005% CHAPS. **(B)** Bacteria recovered from THP-1 macrophage-like cells infected with WT *B. ovis* or a *B. ovis bepE* restored mutant. **(C, D)** Bacteria recovered from infected THP-1 cells treated with 25 µM nicardipine or 25 µM cilnidipine at **(C)** 24- or **(D)** 48-h post infection. Infections were performed three times; values are means ± SD from three independent trials. Statistical significance was calculated at 24 hpi and 48 hpi with an unpaired t-test (*, P < 0.05; **, P < 0.01).

### Restoration of the *B. ovis bepE* reading frame confers resistance to dihydropyridines in the intracellular niche

The enhanced resistance of *bepE*-restored *B. ovis* mutants to drugs and envelope stress in vitro prompted us to examine their impact on bacterial replication and susceptibility to nicardipine and cilnidipine within the host environment. To assess whether *bepE* restoration affects intracellular proliferation, we infected THP-1 macrophage-like cells with either wild-type (WT) *B. ovis* (carrying the *bepE* pseudogene) or a strain with a reconstituted functional *bepE* allele. Although the *bepE*-restored strain showed a modest increase in CFUs at 24 hours, both strains exhibited comparable intracellular burdens at 48 hours post-infection (Fig. 4B).

We next asked whether restoration of the *bepE* open reading frame confers resistance to dihydropyridines in the intracellular niche. Infected macrophages treated with nicardipine or cilnidipine showed a pronounced difference in bacterial burden between strains: the *bepE*-restored mutant exhibited approximately 10-fold higher CFUs at 24 hours and nearly 100-fold higher CFUs at 48 hours compared to WT *B. ovis* (Fig. 4C–D). Despite this substantial increase in drug tolerance, *bepE* restoration did not fully rescue growth under drug treatment to that observed in untreated cells at later time points, suggesting that additional host- or drug-associated factors contribute to the intracellular activity of dihydropyridines. Importantly, reduced bacterial recovery was not due to host cytotoxicity (Fig. S4), consistent with our initial screen. Together, these results demonstrate that restoration of *bepE* enhances *B. ovis* resistance to dihydropyridine Ca^2+^ channel blockers in macrophages, although this protection is partial and depends on the time point post infection.

### *bepE* contributes to bile acid resistance in *B. abortus*

Frameshift mutations in the *B. ovis bepE* pseudogene restored the open reading frame found in other *Brucella* species, conferred resistance to dihydropyridines, and enhanced survival under conditions of cell envelope stress. To determine whether *bepE* similarly influenced chemical resistance in other *Brucella*, we investigated its role in *B. abortus*, a closely related zoonotic pathogen that encodes an intact *bepE* gene (locus tag BAB_RS17475; BAB1_0323). Attempts to delete *bepE* using standard allelic exchange with sacB/sucrose counterselection were unsuccessful. Instead, we inactivated the gene via single-crossover recombination, generating a mutant (Δ*bepE*) that expresses only the first 103 amino acids of BepE. We assessed the sensitivity of WT and Δ*bepE* strains to nicardipine, cilnidipine, and the membrane-disrupting agents deoxycholate and CHAPS (1% w/v) on solid media (Fig. 5A). While 1% deoxycholate modestly impaired WT growth, Δ*bepE* was unable to grow under this condition, consistent with prior findings in *B. suis* [41]. In contrast, *bepE* disruption did not affect sensitivity to CHAPS or the dihydropyridines under the tested conditions.

**Figure 5.**
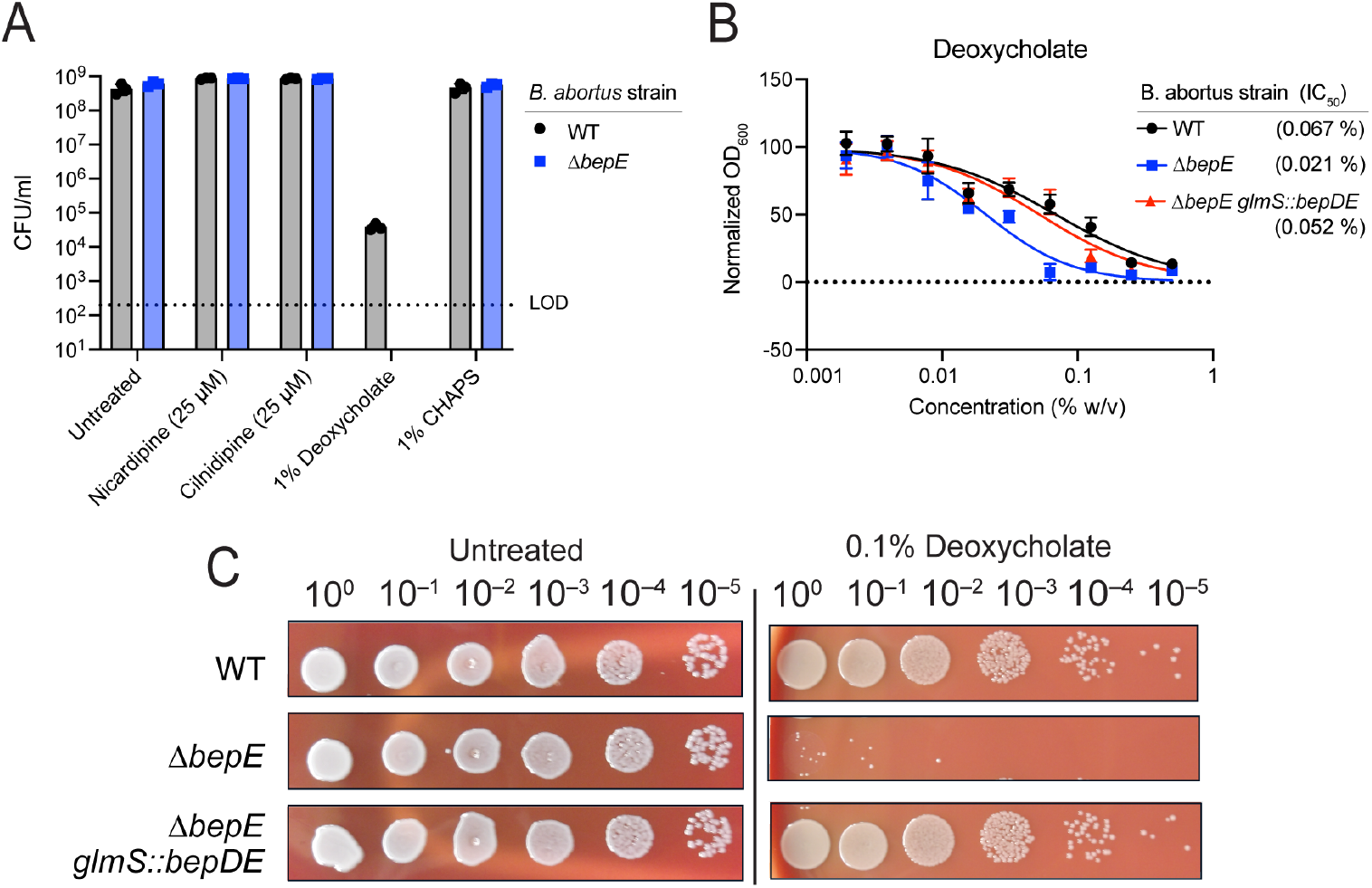
Loss of BepE function renders B. abortus sensitive to deoxycholate. **(A)** Colony-forming units of *B. abortus* WT and Δ*bepE* (BAB_RS17475) enumerated after growth on TSA blood medium alone (Untreated) or containing 25 *µ*M cilnidipine, 25 *µ*M nicardipine, 1% deoxycholate (w/v), or 1% CHAPS (w/v). Dotted line indicates limit of detection. **(B)** Deoxycholate dose-response curves with IC50 values. Growth of *B. abortus* WT, Δ*bepE*, and the complementation strain Δ*bepE glmS::bepDE* in liquid medium with deoxycholate was assessed by optical density after 48 h. Values represent mean ± SD of three independent trials, each normalized to the untreated control. **(C)** Growth of serially-diluted B. abortus strains spotted onto TSA blood plates with or without 0.1% deoxycholate (w/v).

To confirm that the deoxycholate sensitivity of the Δ*bepE* strain was due to loss of BepE function, we integrated a single copy of the *bepDE* operon at the *glmS* locus for genetic complementation. In both liquid and solid media, the Δ*bepE* mutant showed increased sensitivity to deoxycholate (0.1– 1% w/v) and genetic complementation restored resistance to WT levels (Fig. 5B–C). Together, these results demonstrate that BepE specifically contributes to *B. abortus* resistance to a secondary bile acid.

### *bepE* is dispensable for intracellular growth but confers *B. abortus* resistance to cilnidipine treatment in host cells

We assessed whether loss of *bepE* affects *B. abortus* fitness in macrophage infection models. To do this, we used two distinct macrophage systems that differ in their physiological characteristics and potential relevance to *Brucella* infection. THP-1 cells are a well-established model of circulating monocyte-derived macrophages, while fetal liver alveolar-like macrophages (FLAMs) are a primary ex vivo model that more closely resembles tissue-resident macrophages [42]. Because *B. abortus* encounters both monocyte-derived and tissue-resident macrophages during the course of host colonization, we evaluated bacterial replication in both systems to capture this range of possible host cell environments. In THP-1 macrophage-like cells, Δ*bepE* colony-forming units (CFU) recovered at 2, 24, or 48 hours post-infection were not significantly different from those of wild-type or the genetically complemented strain (Δ*bepE glmS::bepDE*) (Fig. 6A). Similarly, in FLAMs, the growth and survival of the Δ*bepE* mutant matched that of the wild-type strain throughout the infection time course (Fig. S5).

**Figure 6.**
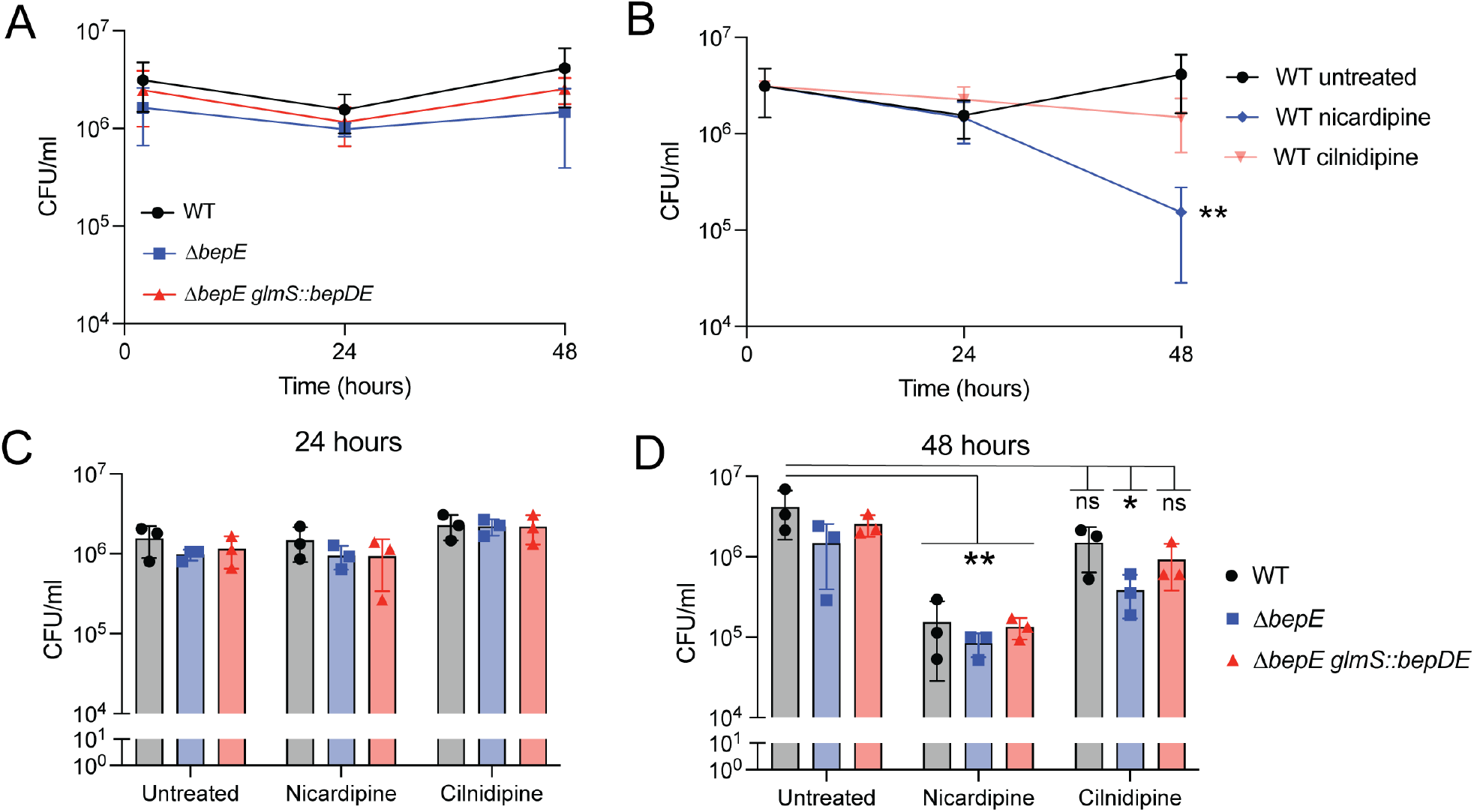
B. *abortus bepE* does not contribute to survival in THP-1 macrophages but confers enhanced resistance to cilnidipine in the intracellular niche. **(A)** *B. abortus* CFU recovered from THP-1 macrophages after infection with WT, Δ*bepE*, and the complementation strain Δ*bepE glmS::bepDE*. **(B)** WT *B. abortus* cells recovered from infected THP-1 macrophages untreated or treated with 25 µM nicardipine or 25 µM cilnidipine. **(C, D)** CFUs of WT, Δ*bepE*, and Δ*bepE glmS::bepDE* recovered from THP-1 macrophages at 24 **(C)** or 48 hpi **(D)**. Infected macrophages were treated with 25 µM nicardipine or 25 µM cilnidipine. Infections were performed three times; values are means ± SD from three independent trials. Statistical significance was assessed at 48 hpi using two-way ANOVA on log10 transformed CFU values, followed by Tukey’s multiple comparison’s test compared with the untreated WT group (*, P < 0.01; **, P < 0.001).

We next tested whether *bepE* influences *B. abortus* sensitivity to dihydropyridines in host cells. Using a luciferase-expressing *B. abortus* strain to monitor intracellular bacteria in THP-1 cells, we found that *B. abortus* was more sensitive to nicardipine (IC_50_ = 4.4 µM) than to cilnidipine (IC_50_ = 13.5 µM) (Fig. S6A). Consistent with this, significantly fewer wild-type *B. abortus* CFUs were recovered from nicardipine-treated THP-1 cells compared to untreated controls (Fig. 6B). Deletion of *bepE* did not alter sensitivity to nicardipine but did increase sensitivity to cilnidipine: the Δ*bepE* strain showed a significant reduction in CFU recovery at 48 hours post-infection relative to wild-type (Fig. 6D), although no difference was observed at 24 hours (Fig. 6C). The complemented strain restored resistance to WT levels (Fig. 6D). These results indicate that *bepE* contributes to *B. abortus* resistance to cilnidipine in the intracellular niche. Importantly, in axenic conditions, both WT and Δ*bepE* B. *abortus* were insensitive to nicardipine and cilnidipine (up to 25 µM) (Fig. S6C-D). We conclude that *B. abortus* is more sensitive to these dihydropyridines in the intracellular environment, owing either to host-directed activity or to increased susceptibility to the drugs within its intracellular niche.

## Discussion

### Dihydropyridines as intracellular pathogen inhibitors

Previous small-molecule screens identified dihydropyridine calcium channel blockers, including nicardipine, as inhibitors of *Brucella abortus* replication in host cells [15, 16]. Through an unbiased small-molecule screen, we show here that treating *B. ovis*-infected macrophage-like cells with micromolar concentrations of nicardipine or cilnidipine significantly reduces bacterial fitness within the intracellular niche. These results extend the anti-infective activity of dihydropyridines to multiple *Brucella* species and macrophage models. Notably, although nifedipine and the non-dihydropyridine calcium channel blocker verapamil have previously demonstrated intracellular anti-mycobacterial or anti-*Brucella* activity [36, 43, 44], neither compound emerged as a hit in our *B. ovis* screen at the concentrations tested.

Beyond *Brucella*, human case control studies have found that the use of dihydropyridine calcium channel blockers is associated with a reduced risk of active tuberculosis [24], suggesting broader clinical potential for this drug class against intracellular pathogens. Indeed, dihydropyridines have been reported to inhibit several intracellular bacteria, including *Mycobacterium tuberculosis* [36], *Coxiella burnetii* [16], and *Legionella pneumophila* [45] highlighting their promise as broad-spectrum intracellular pathogen inhibitors. However, despite these encouraging findings, the cellular and molecular mechanisms by which calcium channel blockers exert their antimicrobial effects (on both host cells and bacteria) remain poorly understood. Some data indicate these drugs may disrupt bacterial efflux systems and potentiate sensitivity to other chemical stressors. For example, the dihydropyridine amlodipine exhibits synergy with imipenem against multidrug-resistant *Acinetobacter baumannii* by inhibiting bacterial efflux [46]. Similarly, the non-dihydropyridine calcium channel blocker verapamil has shown synergistic effects with the anti-tubercular drugs clofazimine and bedaquiline [47]. We have not yet explored potential synergies between dihydropyridines and standard-of-care antibiotics for treatment of brucellosis, but our results raise the possibility that dihydropyridines could be integrated into combination therapies to improve treatment outcomes.

### Nicardipine treatment alters metal homeostasis in macrophages

The effects of calcium channel blockers on THP-1 macrophage physiology and metal homeostasis are not well defined, though L-type Ca^2+^ channels are reported to be expressed in this human cell line [48, 49]. To investigate how nicardipine affects intracellular metal levels in THP-1 macrophage-like cells, we used triple quadrupole inductively coupled plasma mass spectrometry (ICP-QQQ). While most metals remained unchanged, nicardipine treatment caused a significant increase in total intracellular calcium (Fig. 2). Previous studies have shown that blocking voltage-gated Ca^2+^ channels can paradoxically enhance calcium influx in macrophages, leading to upregulation of pro-inflammatory pathways and improved clearance of *M. tuberculosis* [36]. Calcium signaling also influences macrophage polarization, with distinct calcium channel activities promoting either a classically activated (M1) or alternatively activated (M2) state [50]. As *Brucella abortus* has been reported to promote M2 polarization to support its intracellular survival at later infection stages [51], it is possible that increased calcium in nicardipine-treated macrophages may favor M1 polarization and activate antimicrobial pathways.

In addition to increasing intracellular calcium, nicardipine treatment also elevated manganese levels in THP-1 cells (Fig. 2). Manganese (Mn^2+^) is reported to activate antiviral pathways such as STING [38], and plays an important role in antibacterial immunity, including host defense against *M. tuberculosis* [37]. Thus, nicardipine may enhance antimicrobial responses by modulating Mn^2+^ availability, potentially creating a less permissive intracellular environment for *Brucella*. Notably, *Brucella* tightly regulates manganese homeostasis, and perturbation of this balance reduces bacterial fitness both in vitro and in vivo [52].

The specific host pathways and ion channels responsible for the observed calcium and manganese shifts following drug treatment remain undefined. Identifying these systems could reveal new opportunities for host-directed therapeutic strategies. We note that while the ICP-QQQ method we used enables accurate quantification of total intracellular metal levels, it does not resolve how metals are spatially distributed among subcellular compartments or how this distribution changes upon treatment. It is possible that nicardipine alters metal compartmentalization, leading to localized effects on host or pathogen physiology. Despite this limitation, our data show that nicardipine significantly reshapes the macrophage metallome (Fig. 2) without compromising cell viability (Fig. S4), supporting a potential role for host metal modulation in its anti-*Brucella* activity.

### Pseudogenization of bepE underlies B. ovis sensitivity to dihydropyridines and envelope stressors

Although dihydropyridine-induced changes in host metal levels could influence *Brucella* infection dynamics, we found that these compounds also directly inhibit *B. ovis* growth in axenic culture (Fig. 1B & Fig. S3). To investigate the genetic basis of this sensitivity, we selected for spontaneous *B. ovis* mutants capable of growing in the presence of cilnidipine, the more potent of the two dihydropyridines tested. Whole-genome sequencing of resistant isolates revealed two classes of mutations: non-synonymous substitutions in a predicted *emrA*-family efflux gene, and single-nucleotide deletions in *bepE*, a pseudogene encoding a subunit of an RND-family transporter [41, 53]. These deletions restored the *bepE* reading frame, yielding a full-length gene homologous to functional *bepE* loci in other *Brucella* species.

*B. ovis* mutants with restored *bepE* alleles exhibited increased dihydropyridine resistance in the intracellular niche, but they remained partially sensitive to drug at 48 hours post-infection, indicating that BepE-mediated transport processes alone are insufficient to fully protect in host cells. This residual sensitivity suggests that dihydropyridines may also impair *Brucella* fitness through host-directed mechanisms. Consistent with this, we observed that a *B. abortus bepE* mutant was as resistant as wild-type in axenic culture across a range of nicardipine and cilnidipine concentrations (Fig. S6) but became significantly more susceptible to cilnidipine during intracellular infection in THP-1 cells (Fig. 6 & Fig. S6). Notably, wild-type *B. abortus* was also more sensitive to nicardipine than cilnidipine in an intracellular setting (Fig. 6); neither drug inhibited *B. abortus* growth in axenic culture up to 25 µM (Fig. S6). Together, these results support a model in which dihydropyridines act, at least in part, by modulating host pathways that restrict intracellular *Brucella* growth or survival.

The precise mechanism of direct nicardipine and cilnidipine toxicity to *B. ovis* remains undefined, and the lack of resistance mutations outside transport-related genes suggests that cilnidipine may act on essential pathways or exert its effects through multiple mechanisms. *B. ovis* is the only classical *Brucella* species to have lost a functional *bepE* gene (Fig. 3), and our data demonstrate that *bepE* pseudogenization is a key factor in the documented sensitivity of *B. ovis* to chemical stress relative to other *Brucella* spp. [39, 40]. *B. ovis* has undergone extensive genome reduction [19, 54] and we speculate that *bepE* loss of function reflects restriction of this species to the niche of the ram reproductive tract, where exposure to cell envelope stressors may be reduced [55].

### Function and evolutionary flexibility of bepE in the Brucella genus

Although *bepE* contributes to chemical stress resistance, it was dispensable for intracellular survival in two physiologically distinct macrophage models (Fig. 6 and Fig. S5). These findings are consistent with prior work in *B. suis*, where Δ*bepE* mutants replicated normally in cultured epithelial and phagocytic cells. In contrast, Δ*bepC* mutants (which lack a shared outer membrane component used by multiple RND-family efflux systems, including BepE) exhibited reduced spleen colonization in mice [41, 56]. Thus, *bepE* may support fitness when *Brucella* faces specific immune pressures in vivo. Remarkably, single-nucleotide deletions can restore *B. ovis bepE* pseudogene function, highlighting its potential role as a “contingency” locus [57]. Contingency genes are typically non-functional under standard conditions but can be reactivated to confer selective advantages under particular environmental conditions. Such genomic flexibility may allow *Brucella* species to balance the evolutionary pressure for genome streamlining with the need to preserve adaptability. Future studies are necessary to determine whether the presence of a frameshifted *bepE* allele in *B. ovis* reflects selective pressure to maintain a reactivatable transport function or simply represents an early stage in the irreversible degradation of this gene during host restriction.

## Acknowledgements

We thank Rob Abramovitch, Andrew Olive, and Esther Chen for helpful feedback and ideas over the course of this study. We also thank the lab of Andrew Olive for assistance with cultivation of FLAM cells. Research reported in this publication was supported in part by the NIH under award number R01AI177619 to S.C., and P41GM135018 and R01GM038784 to T.V.O.

## Materials and Methods

### Bacterial strains and growth conditions

*Brucella ovis* 25840 and *Brucella abortus* 2308 were grown on tryptic soy agar (TSA; Difco Laboratories) plates supplemented with 5% (v/v) sheep blood (Quad Five) or in brucella broth (BB; Difco Laboratories) dissolved in milliQ water for liquid cultures. *B. ovis* and *B. abortus* cells were incubated at 37ºC with 5% (v/v) CO2 supplementation.

All *Escherichia coli* strains were grown in liquid LB or on LB solidified with 1.5% (w/v) agar. Top10 and WM3064 strains were incubated at 37ºC. WM3064 was grown with 30 µM diaminopimelic acid (DAP) supplementation. The growth medium contained 50 µg ml^-1^ kanamycin or 20 µg ml^-1^ chloramphenicol when necessary. Primer, plasmid, and strain information are available in Table S2.

### Plasmid and strain construction

#### (i) Deletion strain construction

To inactivate *bepE* in *B. abortus* (gene ID BAB_RS17475), a 528-bp internal fragment was cloned into pNPTS138-*cat* (a suicide vector that carries a chloramphenicol resistance gene). The resulting plasmid was used to disrupt the target gene by a single crossover insertion. Positive colonies were selected on sheep blood agar (SBA) plates containing 3 µg ml^**−**1^ chloramphenicol. The obtained mutant strains are predicted to produce only the first 103 amino acids of BepE.

#### ii) Complementation strain construction

To build plasmids for genetic complementation, the coding sequences of *bepD* (BAB_RS17470) and *bepE* were PCR amplified, starting ∼300 bp upstream and ending ∼50 bp downstream of the *bepD/E* operon. The PCR product was purified and inserted into plasmid pUC18-mTn7 by restriction enzyme digestion and ligation, followed by chemical transformation into *E. coli* TOP10 cells. After sequence confirmation by Sanger sequencing, the resulting pUC18-mTn7-*bepD/E* plasmid was transformed into chemically competent *E. coli* WM3064. The plasmid was co-conjugated into *B. abortus* strains with pTNS3, a suicide helper plasmid expressing the *Tn7* integrase gene. *B. abortus* colonies carrying the integrated mTn7-*bepD/E* construct at the *glmS* locus were selected on TSA blood plates containing 50 µg ml^**−**1^ kanamycin.

#### (iii) Bioluminescent strain construction

Luminescent *B. ovis* and *B. abortus* strains were generated from wild-type *B. ovis* ATCC 25840 and *B. abortus* 2308 parent strains by integration of a pUC18-mTn7-gentamicin plasmid harboring the *luxCDABE* operon at the *glmS* locus (the mini-Tn7 plasmid was a gift from H.P. Schweizer). The original gentamicin resistance marker was disrupted by Tth111I restriction enzyme digest (New England Biolabs) and replaced with a kanamycin cassette by Gibson assembly (Table S2). The pUC18-mTn7-*luxCDABE*-kanamycin plasmid was co-conjugated into *B. ovis* and *B. abortus* strains with the suicide helper plasmid pTNS3 expressing the *Tn7* integrase gene, as above for complementation strain construction. Positive *B. ovis* and *B. abortus* colonies carrying the integrated mTn7-*luxCDABE* luminescence construct were selected on TSA blood plates containing 50 µg ml^**−**1^ kanamycin.

### Small-molecule screening

#### (i) Macrophage infection

The drug screen was conducted at the Michigan State University Assay Development and Drug Repurposing Core (ADDRC). Briefly, THP-1 cells were seeded using a liquid handler (BioTek MultiFlo) dispensing 25µl of host cells at a titer of 2 × 10^4^ cells per well with 50 ng mL^**−**1^ phorbol myristate acetate (PMA) to induce differentiation into macrophage-like cells in white 384-well plates (Corning). THP1 cells were incubated at 37ºC in 5% (v/v) CO2 for 72 h. In parallel, luminescent *B*.*ovis* cells were streaked out on TSA blood plates and incubated at 37^0^C in 5% CO2 for 72 hours. Immediately preceding the addition of luminescent *B B. ovis* to differentiated THP-1 cells, screening compounds from the Prestwick Chemical Library dissolved in 2 mM DMSO stock solutions were added to the assay plates by a 150 nanoliter automatic robotic pin transfer (Beckman Coulter Biomek FX^p^) for a final concentration of 5 μM. Then, luminescent *B. ovis* cells were resuspended in RPMI medium supplemented with 10% (w/v) heat-inactivated fetal bovine serum (HI FBS) (HyClone); host cells were infected by transferring 25 µl of luminescent *B. ovis* cell suspension using a liquid handler (BioTek MultiFlo). Plates were centrifuged for 5 min at 150 x *g* and incubated for 1 h at 37ºC in 5% (v/v) CO2. Following incubation, 10 µl of fresh medium containing 300 ug ml^**−**1^ of gentamicin was added to each well and the plate was incubated for 24 hours at 37ºC and 5% (v/v) CO2 for 48 h before evaluating *B. ovis* luminescence using a plate reader (Biotek Synergy Neo Plate Reader). THP-1 cell viability was then determined using a fluorescent Gly-Phe-AFC peptide (MP Biochemicals). Positive hits from this intracellular *B. ovis* screen were defined as small molecules that diminished the *B. ovis* luminescence signal to at least 35% of that for DMSO (vehicle)-treated samples and did not affect THP-1 viability. A Z’ factor of 0.417 was achieved for intracellular inhibitory screening.

#### (ii) Axenic growth

Luminescent *B. ovis* cells were resuspended from a 72-h-old plate in BB (Difco Laboratories), and 60 µl of cells were dispensed into each well of 384-well plates by liquid handling (Biotek MultiFlo). Small molecules were then transferred from compound screening libraries using automated robotic pin transfer (Beckman Coulter Biomek FX). Assay plates were then incubated at 37ºC and 5% (v/v) CO2 for 48 h before evaluating *B. ovis* luminescence on a plate reader (Biotek Synergy Neo Plate Reader). A Z’ factor of 0.473 was achieved for axenic inhibitory screening.

### Growth curves

BB was inoculated with *Brucella* cells from 48-to approximately 72-h-old TSA blood plates at cell densities ranging from OD600 = 0.05 to OD600 = 0.1. Growth was assessed by measuring OD600 on a spectrophotometer (Genesys 30 Visible Spectrophotometer). Growth curves were conducted at least three independent times with two technical replicates in each experiment. Representative curves are shown for each set of strains. Where indicated, 25 µM nicardipine (Cayman Chemical) and 25 µM cilnidipine (Cayman Chemical), dissolved fresh in DMSO (Thermo Scientific), were added to BB at the start of the growth experiment.

### Dose-response growth assay in axenic culture

Two-fold serial dilutions of compounds were prepared manually in 96-well plates, with final concentrations of each compound ranging from 0.2 µM to 50 µM suspended in 100 µl BB. From a 48-h-old TSA blood plate, *B. ovis* or *B. abortus* cells were resuspended in BB at densities ranging from OD600 = 0.05 to OD600 = 0.1. *Brucella* cell suspensions were transferred into 96-well plates (100 µl per well) containing compound dilutions for a final volume of 200 µl. Plates were incubated without shaking at 37ºC and 5% (v/v) CO2 for 48 h; absorbance was read spectrophotometrically at OD600 with a plate reader (Tecan Spark 10M Multimodal Plate Reader or Tecan Infinite M200 Pro microplate reader).

### Forward genetic selection for cilnidipine-resistant B. ovis mutants

To identify spontaneous cilnidipine-resistant mutants, BB was inoculated with wild-type *B. ovis* ATCC 25840 at a final OD600 of ∼0.2 and then plated onto TSA blood plates containing 25 µM cilnidipine and incubated for 72 h at 37ºC and 5% (v/v) CO2. Six independent spontaneous mutants that acquired the ability to grow on cilnidipine-containing TSA blood plates were obtained. These mutant isolates were restreaked onto TSA blood cilnidipine-containing plates, and individual colonies were picked and grown in BB containing 25 µM cilnidipine at 37ºC and 5% (v/v) CO2 to confirm resistance to cilnidipine.

### DNA extraction, amplification, and quantification

Genomic DNA was extracted following a standard guanidium thiocyanate protocol. Cilnidipine-resistant mutants were streaked onto TSA blood plates; individual colonies were picked and cultivated in BB overnight. One milliliter of culture was centrifuged at 12,000 rpm for 20 s at room temperature; the resulting pellet was washed with 0.5 ml phosphate-buffered saline (PBS), pH 7.4. The pellet was resuspended in 0.1 ml TE buffer, pH 8.0 (10 mM Tris-HCl, pH 8.0 [Fisher Bioreagents], 1 mM EDTA pH 8.0 [Fisher Bioreagents]), to which 0.5 ml of 0.5% (v/v) sarkosyl was added. Following a 15-min incubation at 60ºC, 0.25 ml of cold 7.5 M ammonium acetate (Fisher Bioreagents) was added. After incubation for 10 min on ice, 0.5 ml of chloroform (Fisher Bioreagents) was added, and samples were vortexed and centrifuged at 13,000 x *g* for 30 minutes at 4 ºC. The aqueous phase was mixed with 0.54 volume of cold isopropanol (Fisher bioreagents) and incubated at room temperature for 15 min before centrifugation at 13,000 x *g* for 10 minutes. The pellet was washed three times in 70% (v/v) ethanol before being resuspended in TE buffer plus RNase A, at which point the DNA concentration was determined spectrophotometrically (Thermo Scientific NanoDrop One/One^C^ Microvolume UV-Vis Spectrophotometer).

### Whole-genome DNA sequencing

Genomic DNA (gDNA) from the parent *B. ovis* strain and gDNA from six independent spontaneous mutants that grew on TSA blood plates containing 25 µM cilnidipine were purified using the standard guanidium thiocyanate-based procedure described above. DNA library preparation and sequencing were performed at SeqCenter (https://www.seqcenter.com). Reads were mapped to the *Brucella ovis* ATCC 25840 genome (chromosome 1 and chromosome 2 RefSeq accession numbers NC_009505 and NC_009504, respectively), and polymorphisms were identified using breseq [58] (https://github.com/barricklab/breseq).

### Cell culture

THP-1 macrophage-like cells were grown to a maximum titer of 1 × 10^6^ cells ml^**−**1^ in complete RPMI 1640 medium supplemented with 2 mM glutamine (GIBCO), and 10% (v/v) HI FBS (HyClone). Fetal liver alveolar macrophages (FLAM) were prepared as reported by Thomas et al. [59]. Briefly, fetal liver cells were harvested as described [60] and grown in complete RPMI 1640 medium supplemented with 2 mM glutamine (GIBCO), 10% (v/v) HI FBS (HyClone), 30 ng ml^**−**1^ recombinant mouse GM-CSF (PeproTech), and 20 ng ml^**−**1^ recombinant human TGF-β1 (PeproTech). At 70– 90% confluency, cells were lifted using PBS with 10 mM EDTA at 37°C for 10 minutes, followed by gentle scraping. After 1 week, adherent cells acquired a round, alveolar macrophage–like morphology, at which point stocks were frozen for infection studies.

### Macrophage infections

#### i) THP-1 infection

THP-1 cells were seeded at a titer of 1 × 10^5^ cells per well in 96-well plates, and phorbol myristate acetate (PMA) was added at a final concentration of 50 ng µl^**−**1^ to induce differentiation into macrophage-like cells for 72 h prior to infection. *B. ovis* and *B. abortus* cells were resuspended from a 48-h-old plate in RPMI supplemented with 10% (v/v) HI FBS and added to tissue culture plates on the day of infection at a multiplicity of infection (MOI) of 100. Plates were centrifuged for 5 min at 150 x *g* and incubated for 1 h at 37ºC in 5% (v/v) CO2. The medium was removed, and fresh medium containing 50 µg ml^**−**1^ gentamicin was supplied, followed by a 1-h incubation. Following gentamicin incubation, the medium was removed and replaced with fresh medium containing gentamicin and/or 25 µM nicardipine or 25 µM cilnidipine. For the enumeration of colony-forming units (CFUs), THP-1 cells were washed once with PBS, pH 7.4 and then lysed in H2O for 10 min at room temperature at 2 h, 24 h, and 48 h post infection. Lysates were serially diluted, spotted onto TSA blood plates, and incubated at 37ºC in 5% (v/v) CO2 for 48 h to enumerate CFUs.

#### ii) FLAM infection

FLAM cells were seeded at a titer of 5 × 10^4^ cells per well in 96-well plates 1 day prior to infection. *B. abortus* cells were resuspended from a 48-h-old plate in RPMI supplemented with 10% (v/v) HI FBS and added to tissue culture plates on the day of infection at a MOI of 100. Plates were centrifuged for 5 min at 150 x *g* and incubated for 1 h at 37ºC in 5% (v/v) CO2. The medium was removed, and fresh medium containing 50 µg ml^**−**1^ gentamicin was supplied followed by a 1-h incubation. The medium was removed and replaced with fresh medium containing gentamicin and/or 25 µM nicardipine or 25 µM cilnidipine. For the enumeration of CFUs, FLAM cells were washed once with PBS, pH 7.4 and then lysed with H2O for 10 min at room temperature at 2 h, 24 h, and 48 h post infection. Lysates were serially diluted, spotted onto TSA blood plates, and incubated at 37ºC in 5% CO2 for 48 h to enumerate CFUs.

#### ii) Intracellular dose-response

THP-1 cells were seeded at a titer of 1 × 10^5^ cells per well in 96-well plates, and PMA was added at a final concentration of 50 ng µl^**−**1^ to induce differentiation into macrophage-like cells for 72 h prior to infection. *B. ovis* and *B. abortus* cells expressing the *lux* operon were resuspended from a 48-h-old plate in RPMI supplemented with 10% (v/v) HI FBS and added to tissue culture plates on the day of infection at a MOI of 100. The medium was removed, and fresh medium containing 50 µg ml^**−**1^ gentamicin was supplied and incubated for 1 h. The medium was removed and replaced with fresh RPMI supplemented with 10% (v/v) FBS and containing 50 µg ml^**−**1^ gentamicin and various concentrations of each calcium channel blocker (nicardipine or cilnidipine). Luminescence units were read at 48 h post infection using a Tecan Infinite M200 Pro microplate reader in the MSU University Research Containment Facility when working with *B. abortus* or on a Tecan Spark microplate reader when working with *B. ovis*. The IC50 values were determined from dose-response plots of log(inhibitor) versus normalized responses using GraphPad Prism software.

### Colorimetric THP1 XTT cell proliferation assay

THP-1 cells were seeded at a titer of 1 × 10^5^ cells per well in RPMI supplemented with 10% (v/v) FBS and containing 50 ng µl^**−**1^ PMA to induce differentiation into macrophage-like cells in 96-well plates (Corning) for 48–72 h. Varying concentrations of compounds were prepared manually and transferred to plates containing THP1 cells with final concentrations of each compound (suspended in RPMI and 10% [v/v] FBS) ranging from 0.2 µM to 50 µM. After incubation, 50 µl XTT (sodium 3′-[1-(phenyl aminocarbonyl)-3,4-tetrazolium]-bis (4-methoxy6-nitro) benzene sulfonic acid hydrate) (Roche) was added to each well and incubated for 4 h at 37ºC and 5% (v/v) CO2. Following incubation, absorbance of the reaction product was measured at 570 nm and 650 nm.

### Agar plate/stress assays

After 2 days of growth on TSA blood plates, *B. ovis* or *B. abortus* cells were collected and resuspended in sterile PBS to an OD600 = Each strain was serially diluted tenfold in PBS. Two to five microliters of each dilution was spotted onto either TSA blood, TSA blood containing either 0.005–1% (w/v) SDS, 0.009–1% (w/v) CHAPS, or 0.009–1% (w/v) deoxycholate. After 3 days of incubation on TSA blood, growth was documented photographically.

### Inductively coupled plasma mass spectrometry

#### i) Macrophage treatment

THP-1 cells were seeded at a titer of 1 × 10^7^ cells in 100-mm dishes with RPMI supplemented with 10% (v/v) HI FBS and containing 50 ng µl^**−**1^ PMA to induce differentiation into macrophage-like cells for 72 h. The medium was then removed and replaced with fresh medium alone or with 25 µM nicardipine and incubated for 48 h at 37ºC in 5% (v/v) CO2.

#### ii) Cell harvesting and ICP-MS sampling

The medium was removed and cells were washed with PBS, pH 7.4, twice. TrypLE Express Enzyme (1x) (Gibco) without phenol red was added to the plates and incubated for 5–10 min. Cells were gently detached from the plate surface and fresh RPMI supplemented with 10% (v/v) HI FBS was added. Cell solutions from the dishes were removed and transferred to new 15-ml or 50-ml tubes. Cells in suspension were gently mixed and transferred to 15-ml metal-free tubes. Gadolinium-DOTA (Gd-DOTA) MRI contrast agent (40 µM final concentration) was spiked into each 15-ml metal-free tube before centrifugation at 400 x *g* for 3 min at 4 ºC. After the cells were pelleted, 0.4 ml of each supernatant was transferred into a new 15-ml metal-free tube, then the remaining supernatants were removed. Gd-DOTA, a membrane-impermeable complex, remains confined to the extracellular space [61]. By spiking samples with Gd-DOTA, we can quantify the residual media volume in an isolated cell pellet, enabling accurate determination of intracellular element contents by subtracting extracellular elements in the retained media. Furthermore, this ICP-MS sample preparation method minimizes disruptions to metal homeostasis by eliminating washing steps and maintaining cells in their original culture media during sampling.

The resulting tubes were dried at 70 ºC overnight. Concentrated nitric acid (67–69%, 0.18 ml) was added to the tubes, which were then incubated at 70 ºC overnight until the pellet was completely acid-digested. Following the acid-digestion, Milli-Q water was added to the tubes to yield a 3% (v/v) nitric acid matrix. The elemental concentration of all digested samples and calibration standard were determined on a parts per billion (ppb, µg/L) scale using an Agilent 8900 Triple Quadrupole ICP-MS (Agilent) equipped with an Agilent SPS 4 Autosampler, integrated sample introduction system (ISIS), x-lens, and micromist nebulizer. Instrument tuning and ICP-MS calibration stand preparation were performed as previously described [62]. The isotopes selected for analysis of samples were ^31^P, ^23^Na, ^24^Mg, ^32^S, ^39^K, ^44^Ca, ^51^V, ^52^Cr, ^55^Mn, ^57^Fe, ^59^Co, ^60^Ni, ^63^Cu, ^66^Zn, ^75^As, ^80^Se, ^111^Cd, and ^157^Gd.

