## Supplemental Tables and Figures for "Reversion of a RND transporter pseudogene uncovers latent stress resistance in *Brucella ovis*"

**Table S1.** Compound hits identified from high-throughput screening of the Prestwick Chemical library. These compounds passed the initial screening criteria as selectively inhibitory to *B. ovis* during THP-1 macrophage infection with limited axenic activity.

| Sample | Compound name | Therapeutic class | Major pathway or target |
|---|---|---|---|
| MSU-51 | Trichlorfon | Pesticide | Acetylcholinesterase inhibitor |
| MSU-339 | Guanfacine hydrochloride | Hypertension | Adrenergic receptor agonist |
| MSU-270 | Fendiline hydrochloride | Antiarrhythmic | Calcium |
| MSU-368 | Bepridil hydrochloride | Angina | Calcium |
| MSU-383 | Nicardipine hydrochloride | Hypertension | Calcium |
| MSU-1264 | Lomerizine hydrochloride | Migraine | Calcium |
| MSU-1188 | Cilnidipine | Hypertension | Calcium |
| MSU-1180 | Bifonazole | Antimicrobial | Cell membrane (fungi) |
| MSU-1011 | Flucloxacillin sodium | Antimicrobial | Cell wall |
| MSU-470 | Ceforanide | Antimicrobial | Cell wall |
| MSU-700 | Cefmetazole sodium salt | Antimicrobial | Cell wall |
| MSU-1212 | Ezetimibe | Cholesterol absorption | Cholesterol |
| MSU-370 | Benzbromarone | Gout | Cytochrome P450 |
| MSU-605 | Carbadox | Antimicrobial | DNA synthesis |
| MSU-350 | Clozapine | Antipsychotic | Dopamine |
| MSU-360 | Droperidol | Nausea | Dopamine |
| MSU-374 | Methylergometrine maleate | Uterine atony | Dopamine |
| MSU-980 | Piribedil hydrochloride | Depression | Dopamine |
| MSU-1163 | Aripiprazole | Antipsychotic | Dopamine, serotonin |
| MSU-626 | Racecadotril | Diarrhea | Enkephalinase |
| MSU-976 | Tracazolate hydrochloride | Sedative | GABA |
| MSU-457 | Meclozine dihydrochloride | Nausea | Histamine |
| MSU-589 | Azelastine HCl | Allergic conjunctivitis | Histamine |
| MSU-888 | Promethazine hydrochloride | Allergy | Histamine |
| MSU-1260 | Ritonavir | HIV | HIV protease |
| MSU-973 | Pirlindole mesylate | Depression | Monoamine oxidase |
| MSU-173 | Tranylcypromine hydrochloride | Depression | Monoamine oxidase |
| MSU-144 | Loperamide hydrochloride | Diarrhea | Mu-opioid receptors |

| | | | |
|---|---|---|---|
| MSU-581 | Reboxetine mesylate | Depression | Noradrenaline |
| MSU-1211 | Ipriflavone | Osteoporosis | Osteoclast |
| MSU-376 | Clofazimine | Leprosy | Peroxisome |
| MSU-587 | Cilostazol | Vasodilator | Phosphodiesterase |
| MSU-977 | Zardaverine | Cancer | Phosphodiesterase |
| MSU-142 | Dipyridamole | Anticoagulant | Phosphodiesterase |
| MSU-1031 | Halofantrine hydrochloride | Malaria | Porphyrin |
| MSU-533 | Phenacetin | Analgesia | Prostaglandin |
| MSU-660 | Avermectin B1a | Antimicrobial | Protein synthesis |
| MSU-476 | Primaquine diphosphate | Malaria | Reactive oxygen species |
| MSU-1105 | Verteporfin | Macular degeneration | Reactive oxygen species |
| MSU-590 | Etretinate | Psoriasis | Retinoic acid |
| MSU-531 | Pirenperone | Anxiety | Serotonin |
| MSU-979 | Ozagrel hydrochloride | Thrombosis | Thromboxane |
| MSU-853 | Liothyronine | Hypothyroidism | Thyroid |
| MSU-494 | Propylthiouracil | Hyperthyroidism | Thyroid peroxidase |
| MSU-381 | Lidoflazine | Experimental | Unknown |
| MSU-421 | Suloctidil | Experimental | Unknown |
| MSU-550 | Parthenolide | Dermatitis | Unknown |
| MSU-1013 | Deptropine citrate | Experimental | Unknown |
| MSU-1054 | Levopropoxyphene napsylate | Cough | Unknown |

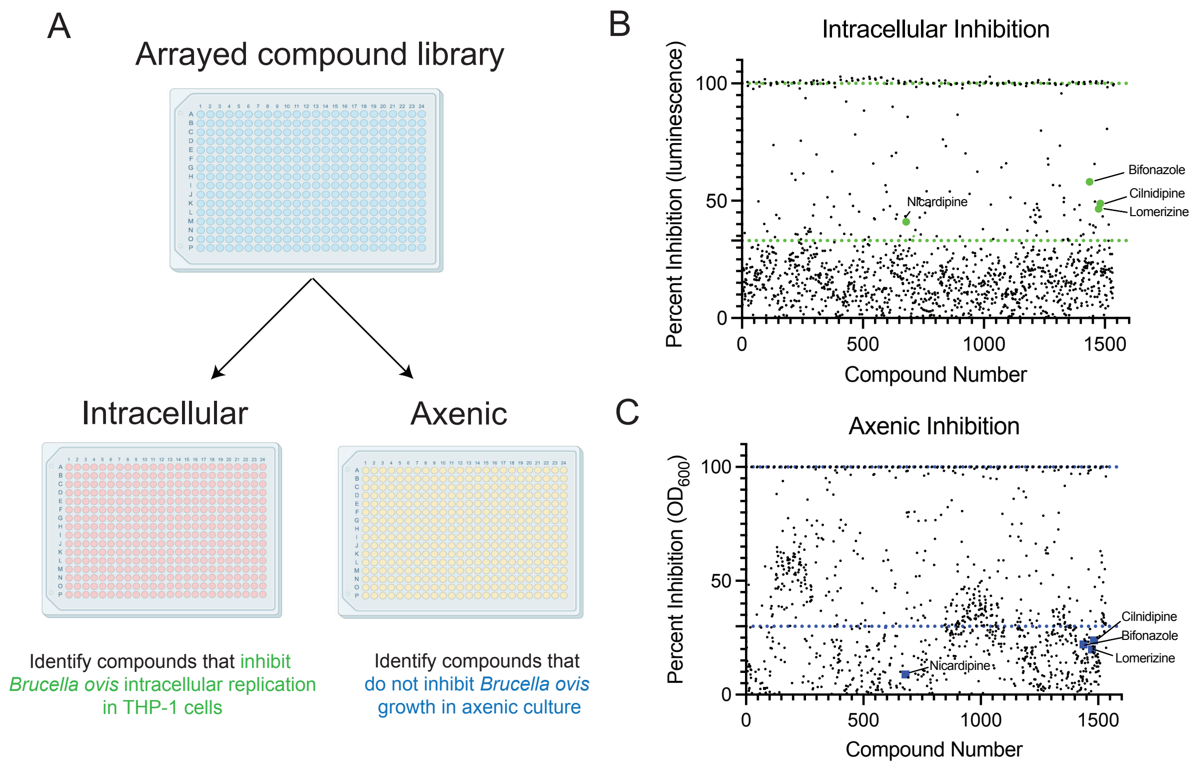


**Figure S1.** Complementary small-molecule screens identify compounds that selectively inhibit *B. ovis* intracellular growth in THP-1 macrophage-like cells. (A) Diagram of the drug screening pipeline for the identification of small molecules that selectively inhibit *B. ovis* intracellular growth with minimal axenic activity. (B) Intracellular inhibition of all tested small molecules, shown as percentage of luminescence emitted by *B. ovis* cells harboring the *lux* operon. Highlighted in green are drug candidates that inhibited *B. ovis* intracellular growth in THP-1 macrophages. The screening of the Prestwick Chemical library had a Z' factor of 0.417 for inhibition of intracellular growth. Dotted line represents hit determination of 35% intracellular inhibition. (C) Effect of small molecules on *B. ovis* growth inhibition in axenic culture. The screening of the Prestwick Chemical library had a Z' factor of 0.473 for axenic growth inhibition. Dotted line represents 35% axenic inhibition. The compounds highlighted in green in B and blue in C are hits that inhibited *B. ovis* intracellular growth but had minimal axenic activity based on our screening criteria.

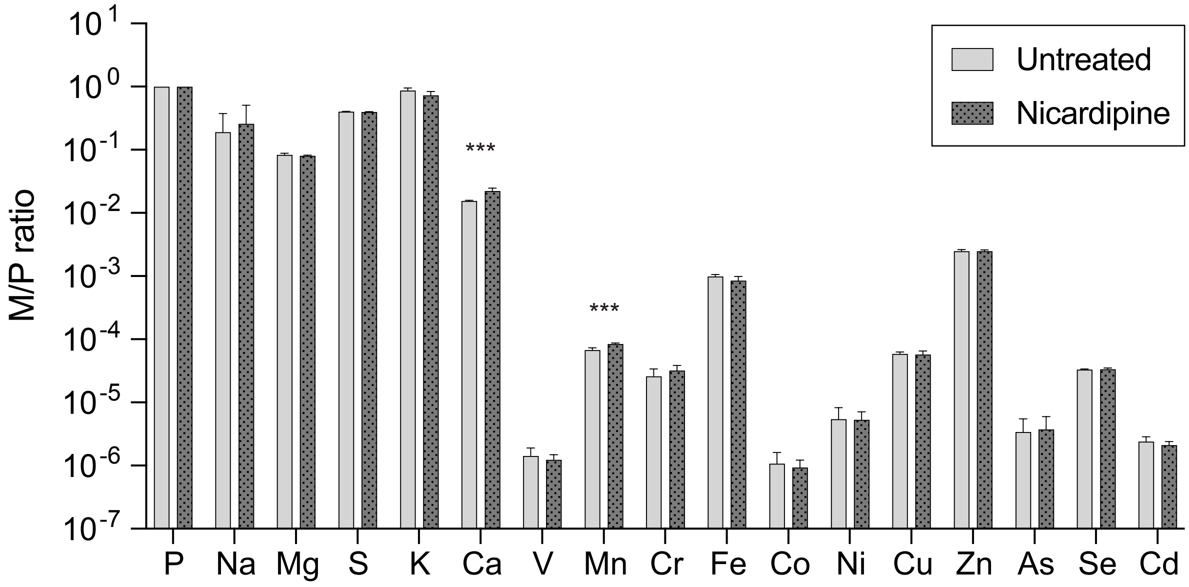


**Figure S2.** Full elemental profile of THP-1 untreated or treated with 25 µM nicardipine for 48 hours. Element content was determined by triple quadrupole inductively coupled plasma mass spectrometry (ICP-QQQ). Levels of each element were normalized to total phosphorus levels (M/P). Bars represent the mean ± standard deviation of 7 biological replicates measured over 2 independent experiments. The M/P ratios for each metal were compared using multiple unpaired t-tests and the Bonferroni-Dunn method to adjust for multiple comparisons (***, adjusted $P < 0.001$).

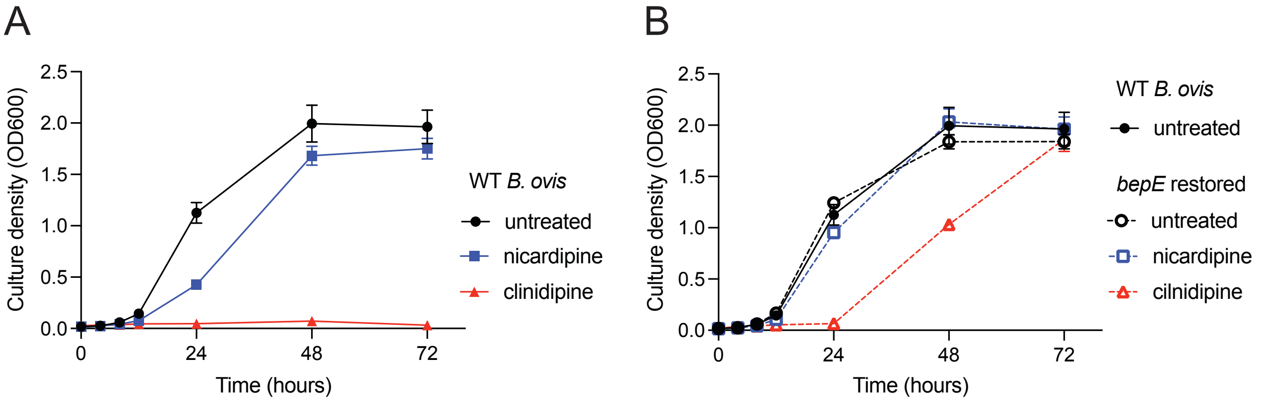


**Figure S3.** Restoration of *B. ovis bepE* confers resistance to dihydropyridine calcium channel blockers during growth in liquid culture. (A) Growth of wild-type (WT) *B. ovis* cultures, untreated (black) 25 $\mu$M nicardipine (blue) or 25 $\mu$M cilnidipine (red), was monitored by optical density at 600 nm. (B) Growth of the *bepE* restored strain in the same treatments as in (A). The WT untreated culture is presented in both panels for reference.

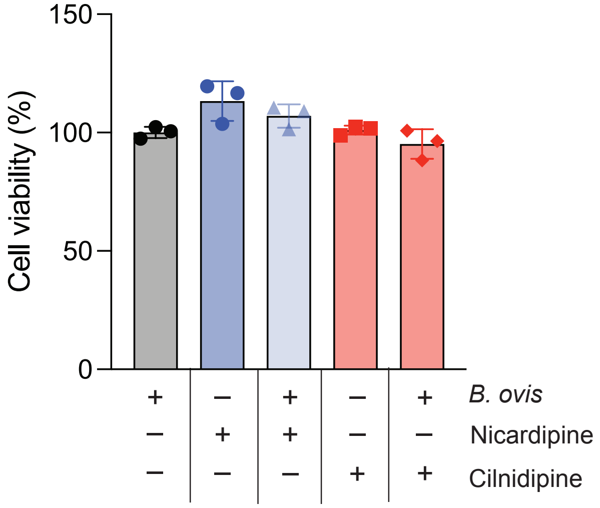


**Figure S4.** *B. ovis* infection and treatment with calcium channel blockers does not affect THP-1 host cell viability. Viability of THP-1 cells, assessed at 48 h post infection or following treatment with 25 $\mu$M nicardipine or 25 $\mu$M cilnidipine with the XTT-cell proliferation assay. Viability was normalized to wells containing untreated THP-1 cells infected with *B. ovis* and to wells only containing cell culture medium (blank). Values are means ± SD from three independent trials.

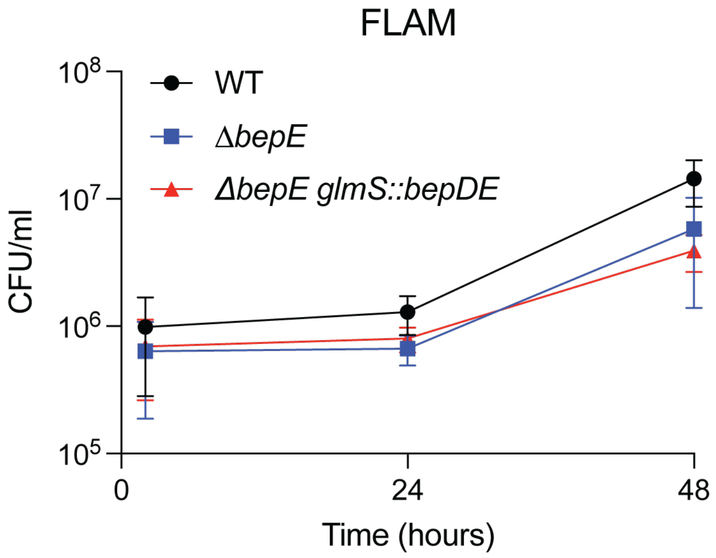


**Figure S5.** *bepE* does not contribute to *B. abortus* survival in fetal liver alveolar macrophages (FLAM). *B. abortus* (WT, Δ*bepE*, and the complementation strain Δ*bepE glmS::bepDE*) recovered from FLAM cells after infection. Values are means ± SD CFU recovered from three independent trials.

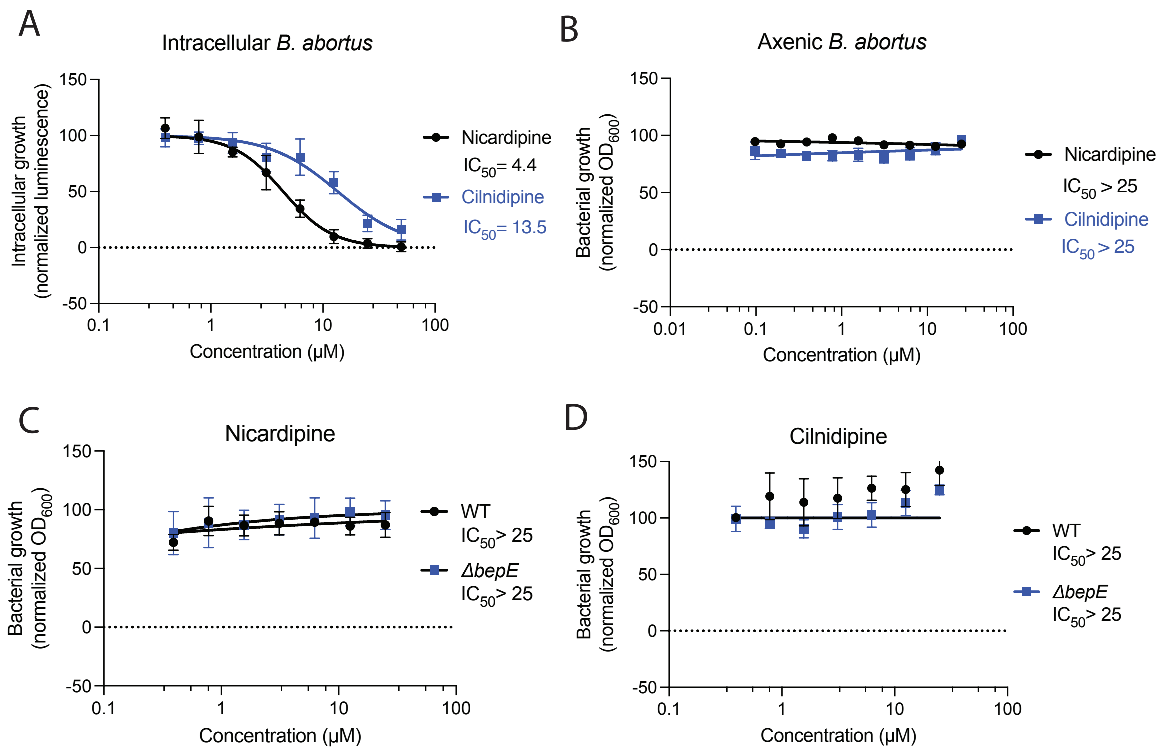


**Figure S6.** Nicardipine and cilnidipine are potent inhibitors of intracellular, but not axenic growth of *B. abortus*. (A) Intracellular inhibitory activities of nicardipine or cilnidipine during infection of THP-1 macrophages by *lux* expressing *B. abortus*. Luminescence was measured after 48 h and normalized to untreated infected controls. (B) Axenic inhibitory activity of nicardipine and cilnidipine during *B. abortus* WT growth in liquid medium. Optical density at 600 nm was measured at 48 h and normalized to untreated cultures. (C, D) Disruption of *bepE* does not affect axenic sensitivity of *B. abortus* to nicardipine or cilnidipine. Growth was measured and analyzed as in panel B.
